# Structured Sampling of Molecularly Classified Mossy Fiber Inputs by Cerebellar Granule Cells

**DOI:** 10.1101/2025.10.02.679866

**Authors:** Xiaomeng Han, Elif Sevde Meral, Jeff Lichtman

## Abstract

The cerebellar granule cell layer receives mossy fiber inputs from diverse brain regions, yet the principles governing how individual granule cells sample distinct types of inputs remain poorly understood. Using a volumetric correlated light and electron microscopy (vCLEM) dataset from an adult female mouse cerebellum, in which VGluT1-positive and VGluT1-negative mossy fiber terminals are molecularly distinguished, we reconstructed granule cell and mossy fiber connectivity to examine input selection rules. We constructed spatially constrained null models to simulate sampling during adulthood and development. Granule cell-centered analysis showed that granule cells shared less innervation from the same mossy fiber than expected by chance. Moreover, subpopulations of granule cells preferentially sample either VGluT1-positive or VGluT1-negative mossy fibers. In contrast, mossy fiber–centered analysis showed that individual terminals distributed their outputs across granule cells in a pattern consistent with random sampling. However, sampling in the adult state was more selective than in developmental simulations. Together, our findings demonstrated structured, non-random sampling of cerebellar VGluT1-positive and VGluT1-negative mossy fiber inputs and provide a framework for understanding how granule cells integrate molecularly distinct inputs to support cerebellar computation.

## Introduction

The cerebellar granule cell layer is the most densely packed neuronal structure in the brain and serves as the first major site of synaptic integration in the cerebellar cortex (Voogd and Glickstein 1998; Eccles 2013). Granule cells receive inputs from extrinsic mossy fibers—axons originating from diverse brain regions such as the pontine nuclei, vestibular nuclei, reticular formation, and the spinal cord—that convey sensory, motor, and state-related information (Voogd and Glickstein 1998; Apps and Hawkes 2009). Intrinsic mossy fiber terminals originate from unipolar brush cells (UBCs) in the vestibulocerebellum (Nunzi and Mugnaini 2000). The convergence of these heterogeneous inputs allows granule cells to expand and transform incoming signals, forming the basis for cerebellar computations, including timing, prediction, and learning (Marr 1969; Albus 1971; Chabrol et al. 2015; Billings et al. 2014).

Based on the Marr-Albus theory (Marr 1969; Albus 1971), granule cells sample mossy fibers in a random, unstructured manner to enhance pattern separation. Recent advances in connectomics (Swanson and Lichtman 2016; Helmstaedter 2013) using volumetric electron microscopy (vEM) (Peddie et al. 2022) have enabled detailed reconstruction of neural circuits. Using these approaches, granule cell input sampling has been shown to be not entirely random: individual granule cells share mossy fiber inputs more frequently than expected under anatomically constrained random models, and mossy fiber terminals are represented unevenly across granule cells (Nguyen et al. 2022). These findings are consistent with theoretical predictions that sparse and structured synaptic connectivity can optimize pattern separation and the dimensionality of cerebellum-like circuits, including granule cell networks (Litwin-Kumar et al. 2017; Cayco-Gajic, Clopath, and Silver 2017). Despite the well-characterized anatomical organization of this layer, the principles by which individual granule cells sample from different mossy fiber types remain largely unknown.

Mossy fibers differ not only in their anatomical origin but also in their molecular identity. Expression of vesicular glutamate transporters (VGluTs) (Fremeau et al. 2004) distinguishes mossy fiber classes: many precerebellar neurons, including those in the lateral reticular nucleus, coexpress VGluT1 and VGluT2 and project to granule cells (Li et al. 2020). In the cerebellum vermis, mossy fiber terminals show heterogeneous VGluT expression, with spinocerebellar fibers and dorsal column fibers preferentially expressing VGluT2 and VGluT1, respectively, forming complementary parasagittal bands in the granule cell layer (Gebre, Reeber, and Sillitoe 2012). In the mouse vestibulocerebellum, mossy fiber terminals that originated from calretinin-positive UBCs are positive for VGluT1 and VGluT2, while those originating from mGluR1α-positive UBCs are positive for VGluT1 only (Nunzi, Russo, and Mugnaini 2003). This molecular diversity suggests that granule cells receive inputs with distinct physiological properties and potentially divergent functional roles. Despite these insights, it remains unclear whether granule cells integrate molecularly distinct mossy fiber types through structured or stochastic sampling.

Advances in large-scale volume electron microscopy (vEM)(Peddie et al. 2022) and correlated light and electron microscopy (CLEM)(de Boer, Hoogenboom, and Giepmans 2015) now enable precise reconstruction of neuronal connectivity while simultaneously capturing molecular identity. This capability is particularly powerful for studying cerebellar granule cell inputs, where mossy fibers differ not only anatomically but also in VGluT expression. Here, we leverage a volumetric CLEM (vCLEM) dataset from the adult mouse cerebellum (Han et al. 2024) in which VGluT1-positive and VGluT1-negative mossy fiber terminals are molecularly distinguished (Fig. 1, a). Using this dataset, we reconstructed granule cells and their mossy fiber inputs and developed spatially constrained null models to simulate input sampling under distinct anatomical constraints.

**Figure 1.**
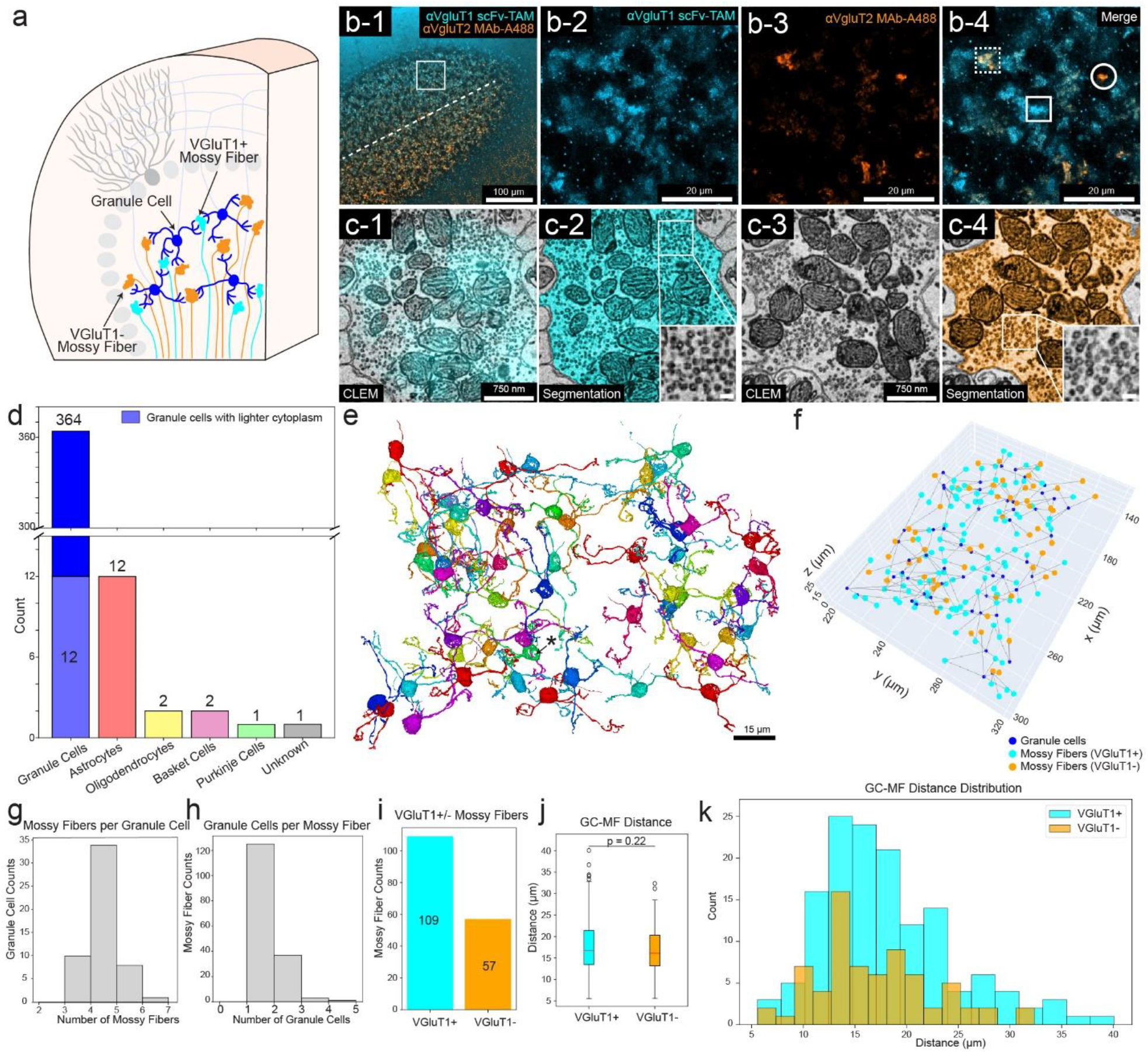
Granule cell layer composition and connectivity with VGluT1+ and VGluT1– mossy fiber terminals. (**a**) Schematic of granule cells and their dendritic claws (blue) connecting VGluT1+ (cyan) and VGluT1– (orange) mossy fiber (MF) terminals in the cerebellar granule layer in the published vCLEM dataset (Han et al. 2024). (**b-1**) Immunolabeling of VGluT1 (cyan) and VGluT2 (orange) in cerebellar hemisphere Crus I, showing complementary expression across the midline (dashed line). (**b-2**–**b-4**) Enlarged single- and dual-channel images from the boxed region in (**b-1**). Solid box: VGluT1+ MF terminals; dashed box: VGluT1+/VGluT2+ MF terminals; circle: VGluT2+ MF terminals. VGluT1+ boutons include both VGluT1+/VGluT2+ and VGluT1+/VGluT2–, while VGluT1– boutons are VGluT2+ only. VGluT1+ terminals are much more abundant than VGluT1-terminals. Reused from (Han et al. 2024) with the authors’ permission. (**c-1**–**c-4**) CLEM images of individual MF terminals from the published dataset (Han et al., 2024). (**c-1, c-2**) VGluT1+ MF terminals identified by overlapping VGluT1 immunolabeling fluorescence (cyan), segmented in cyan, with inset showing synaptic vesicles. (**c-3, c-4**) VGluT1– MF terminals lacking overlapping VGluT1 immunolabeling fluorescence, segmented in orange, with inset showing synaptic vesicles. (**d**) Cell type counts from the reconstructed volume: 364 granule cells (blue), 12 astrocytes, 2 oligodendrocytes, 2 basket cells, 1 Purkinje cell, and 1 unidentified cell. A subset of 12 granule cells displayed lighter cytoplasm. (**e**) 3D reconstructions of 53 granule cells fully contained in the dataset, each shown in a distinct color. Asterisks mark unlabeled regions in the automated segmentation. (**f**) (d) Spatial map of center-of-mass coordinates of 53 granule cells (blue) and their 166 presynaptic MF terminals (VGluT1+: cyan; VGluT1–: orange). Black lines connect synaptically paired GC–MF partners. Interactive version in Supplementary Materials. (**g, h**) Distributions of (**g**) number of MFs per granule cell and (**h**) number of granule cells per MF. (**i**) Counts of VGluT1+ and VGluT1– boutons among the 166 reconstructed MFs. (**j**) Distances between GC–MF partners for VGluT1+ vs. VGluT1– boutons. Welch’s t-test, p = 0.22. (**k**) Distribution of GC–MF distances for VGluT1+ and VGluT1– boutons.

To capture both mature connectivity and the potential developmental origins of granule cell input structure, we modeled two conditions: adult spatial constraints, based on the average dendritic length of granule cells, and developmental potential, inferred from the maximum dendritic reach, reflecting evidence that granule cells exhibit more and longer dendrites during early postnatal development (Dhar, Hantman, and Nishiyama 2018; Altman 1972; Powell et al. 1997). We analyzed input sampling patterns using both granule cell–centered and mossy fiber–centered frameworks to assess whether either side of the synapse demonstrates non-random wiring behavior.

Our results reveal that granule cells in the adult cerebellum sample mossy fiber inputs in a significantly more structured manner than predicted by chance. Subpopulations of granule cells preferentially sampled either VGluT1-positive or VGluT1-negative terminals. In contrast, mossy fiber terminals distributed their outputs to granule cells in a manner largely consistent with random sampling, although with subtle input-type–specific differences. These findings demonstrate that the integration of molecularly distinct mossy fiber inputs by granule cells is not random, but shaped by both spatial and developmental constraints, providing new insight into the structural rules governing cerebellar microcircuit computation.

## Materials and Methods

### vCLEM Dataset

All analyses were performed on a previously published volumetric correlated light and electron microscopy (vCLEM) dataset from the Crus I region of the cerebellar hemisphere of an adult female mouse (Han et al. 2024). In this dataset, mossy fiber terminals were molecularly distinguished using anti-VGluT1 single-chain variable fragment (scFv) immunolabeling, enabling classification of boutons as VGluT1-positive or VGluT1-negative. The dataset was acquired using correlated confocal fluorescence imaging and serial-section scanning electron microscopy, followed by automated stitching, alignment, and 3D segmentation with manual proofreading. Full details of dataset acquisition and processing are provided in (Han et al. 2024).

### Cell Reconstruction and Classification

A total of 382 cells were reconstructed within the middle plane of the dataset by manually assembling segments assigned to the same cell from the automated 3D segmentation. Cell types were classified according to established ultrastructural features (Palay and Chan-Palay 2012). Of these, 364 were identified as granule cells, and the remainder included astrocytes, oligodendrocytes, basket cells, one Purkinje cell, and one cell with unknown identity. For connectivity analysis, we use the subset of 53 granule cells whose dendritic claws terminated exclusively on mossy fiber terminals fully contained within the image volume. Mossy fiber terminals were classified as VGluT1-positive or VGluT1-negative according to immunolabeling in the vCLEM dataset.

### Granule Cell-Mossy Fiber Terminal Connectivity Analysis

Granule cell–mossy fiber synaptic connections were identified by two ultrastructural features: (1) dendritic claws of granule cells encircling mossy fiber terminals, and (2) synaptic contacts characterized by clustered vesicles at the presynaptic terminal in electron microscopy images.

Center-of-mass coordinates were manually annotated for both granule cell soma and mossy fiber terminals. Distances between connected pairs were calculated in 3D. Summary statistics include mean, standard deviation, and maximum distances.

### Null Models

Two spatially constrained null models were implemented. In the adult model, a sphere with a radius equal to the average granule cell dendritic length (17.62 μm) was centered on each cell, and connections were randomly assigned to mossy fibers whose centers of mass fell within the sphere. The number of connections was fixed to match empirical data. In the developmental model, the sphere radius was set to the maximum observed dendritic length (40.08 μm). For mossy fiber–centered analyses, spheres of equivalent radii were centered on mossy fiber terminals. Each model was run with 1,000 Monte Carlo simulations.

### Statistical Analysis

Distances between VGluT1-positive and VGluT1-negative GC–MF pairs were compared using Welch’s t-test. For comparisons between empirical measurements and null-model simulations, one-sided empirical p-values were computed from 1,000 Monte Carlo iterations. Specifically:

- When the empirical statistic was lower than the null mean (e.g., fraction of GC–GC pairs sharing ≥1 MF):

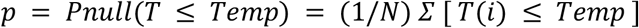

implemented in code as np.mean(null_stats <= emp_stat).
- When the empirical statistic was higher than the null mean (e.g., fraction of GC subtype targets where real > null):

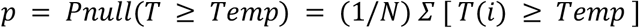

implemented in code as np.mean(null_stats >= emp_stat).

Effect sizes were quantified as Cohen’s d relative to the null distribution, calculated as:

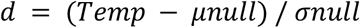

Differences in input fractions were assessed using Mann–Whitney U tests. All analyses and visualizations were performed in Python. The complete Jupyter Notebook used for these analyses is provided in the Supplementary Materials.

## Results

### Molecular and Cellular Profiling of the Granule Cell Layer

We began by characterizing the molecular features of mossy fiber terminals and cellular composition of the granule cell layer within the published vCLEM dataset from an adult female mouse cerebellum (Crus 1 of the hemisphere) (Han et al. 2024). Prior analysis of this dataset revealed that VGluT1-positive mossy fiber terminals are significantly more abundant and exhibit higher synaptic vesicle density and volume than VGluT1-negative terminals (Han et al. 2024)(Fig. 1, b-1 to b-4, c-1 to c-4).

We reconstructed a total of 382 cells located within the middle plane of the dataset (Sup. Fig. 1, a). Among these, 364 were identified as granule cells, while the remainder included 12 astrocytes, 2 oligodendrocytes, 2 basket cells, 1 Purkinje cell, and 1 cell of unknown identity (Fig. 1, d). Notably, oligodendrocytes were observed to myelinate mossy fiber axons (Sup. Fig. 1, c).

Among the 364 granule cells, a subset of 12 granule cells displayed lighter cytoplasmic contrast in EM images (Fig. 1, d; Sup. Fig. 1, b), though the functional significance of this feature remains unclear. Among the 364 granule cells, we identified 53 in which all dendritic claws terminated on mossy fiber boutons fully contained within the image volume (Fig. 1, e). These 53 cells formed synapses with 166 mossy fiber terminals, including 109 VGluT1-positive and 57 VGluT1-negative boutons (Fig. 1, f and i). Each granule cell contacted an average of 4.00 ± 0.65 (mean ± std) mossy fiber terminals (Fig. 1, g). Conversely, each mossy fiber terminal contacted an average of 1.28 ± 0.52 (mean ± std) granule cells (Fig. 1, h). We measured the spatial distance between the center of mass of each granule cell and its connected mossy fiber terminals. The overall average distance was 17.62 ± 6.22 (mean ± std) μm, with a maximum of 40.08 μm. For VGluT1-positive terminals, the mean distance was 17.98 ± 6.44 (mean ± std) μm (maximum: 40.08 μm) (Fig. 1, j), while VGluT1-negative terminals showed a slightly shorter average distance of 16.91 ± 5.74 (mean ± std) μm (maximum: 32.37 μm) (Fig. 1, j). These values are consistent with prior measurements of granule cell dendritic reach (Nguyen et al. 2022), suggesting that while our sample may favor cells with shorter dendrites due to volume constraints, it remains representative. No significant differences were observed in the distances of VGluT1-positive versus VGluT1-negative terminals (Welch’s t-test, p=0.22; Fig. 1, j and k). The center-of-mass coordinates of this set of 53 fully reconstructed granule cells and their 166 mossy fiber partners (Fig. 1, f) formed the basis for the quantitative modeling and connectivity analyses described below.

### Granule Cell-Centered Analysis Reveals Structured Sampling of Mossy Fiber Inputs

To assess whether granule cells sample mossy fiber inputs randomly or in a structured manner, we built two spatially constrained null models centered on each granule cell (Fig. 2, a). The first model (Fig. 2, b; Adult null model) simulated the adult state by constructing a sphere centered at each granule cell using the average dendrite length (17.62 μm) of the 53 complete cells, and randomly assigned connections to mossy fiber terminals whose centers of mass fell within the sphere. The number of connections was fixed to match the empirical data. The second model (Fig. 2, c; Developmental null model) simulated a developmental state by using the maximum dendrite length (40.08 μm) observed among the 53 granule cells. For both models, we ran 1,000 Monte Carlo simulations.

**Figure 2.**
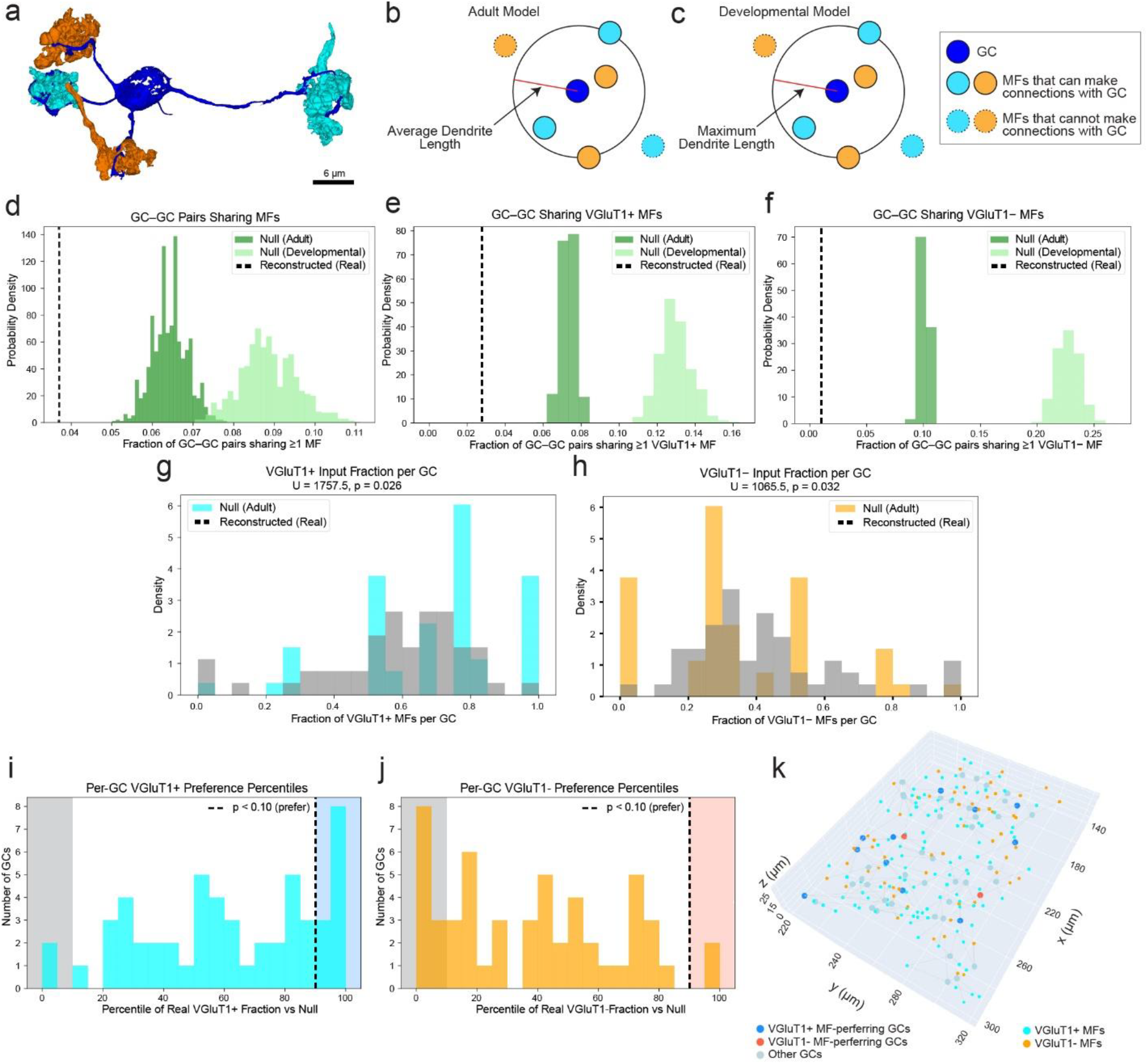
Granule cells exhibit structured, non-random sampling of mossy fiber inputs. (**a**) 3D reconstruction of a centered granule cell (blue) connected to two VGluT1^+^ (cyan) and two VGluT1^-^(orange) mossy fiber terminals. (**b, c**) Null models of GC-centered connectivity. (**b**) Adult null model: connections assigned within a sphere defined by the average dendrite length (17.62 μm). (**c**) Developmental null model: connections assigned within a sphere defined by the maximum dendrite length (40.08 μm). (**d–f**) Probability density distributions of the fraction of GC–GC pairs sharing ≥1 MF. Real data (black dashed line) are compared with null distributions from the adult (dark green) and developmental (light green) models. (**d**) All MFs, (**e**) only VGluT1+ MFs, (**f**) only VGluT1– MFs. (**g**,**h**) Fractions of VGluT1+ (**g**) or VGluT1– (**h**) MF inputs per GC in the real dataset (cyan/orange) compared to the adult null model (gray). Mann–Whitney U test, VGluT1+: U = 1757.5, p = 0.026; VGluT1–: U = 1065.5, p = 0.032. (**i**,**j**) Percentile ranks of each GC’s VGluT1+ (**i**) or VGluT1– (**j**) input fraction relative to the adult null model. (**i**) Granule cells above the 90th percentile (blue shading) were classified as VGluT1^+^-preferring. Granule cells below the 10th percentile (gray shading) correspond to the VGluT1^−^-preferring group shown in (**j**). (**j**) Granule cells above the 90th percentile (blue shading) were classified as VGluT1^−^-preferring, while those below the 10th percentile (gray shading) correspond to the VGluT1+-preferring group in (**i**). (**k**) Spatial map of the three types of GCs (VGluT1+ preferring (blue), VGluT1-preferring (red), and the rest (gray)) and the MF terminals (VGluT1+: cyan; VGluT1–: orange). Black lines connect synaptically paired GC–MF partners. Interactive version in Supplementary Materials.

We first compared the number of granule cell pairs that share at least one mossy fiber input between the empirical data and the null models. The number of shared mossy fiber partners in the real dataset was significantly lower than in both the adult and developmental null models (Fig. 2, d; Adult null model, p-value (real < null) = 0.0000, Cohen’s d = -6.254; Developmental null model, p-value (real < null) = 0.0000, Cohen’s d = -7.408). This suggests that granule cells are not sampling inputs randomly but instead may be structured to maximize access to a broader range of mossy fibers. Furthermore, the deviation (Fig. 2, d, measured by Cohen’d values) was greater in the adult model than in the developmental model, indicating that selective synapse making or pruning from development to adulthood may enhance this structured sampling.

When broken down the mossy fibers into two categories (VGluT1-positive and VGluT1-negative), we observed the number of shared mossy fiber partners in the real dataset was significantly lower than in both the adult and developmental models (Fig. 2, e, VGluT1-positive MFs; Adult null model, p-value (real < null) = 0.0000, Cohen’s d = -12.711; Developmental null model, p-value (real < null) = 0.0000, Cohen’s d = -12.140) (Fig. 2, d, VGluT1-negative MFs; Adult null model, p-value (real < null) = 0.0000, Cohen’s d = -26.482; Developmental null model, p-value (real < null) = 0.0000, Cohen’s d = -21.476) no matter if the mossy fiber terminal is VGluT1-positive or VGluT1-negative. We also observed a stronger deviation from the null models for VGluT1-negative terminals, as quantified by Cohen’s d (Fig. 2, e and f). This supports the hypothesis that granule cells may adopt a strategy to more effectively sample sparser input types such as VGluT1-negative mossy fibers.

We next quantified the fraction of VGluT1-positive and VGluT1-negative terminals contacted by each granule cell. Compared to the adult null model, the real dataset showed an overrepresentation of GCs strongly biased toward VGluT1-positive inputs, as well as a smaller group biased toward VGluT1-negative inputs (For VGluT1-positive MF fraction, Mann–Whitney U test, U = 1757.5, p = 0.026 < 0.05) (For VGluT1-negative MF fraction, Mann–Whitney U test, U = 1065.5, p = 0.032 < 0.05) (Fig. 2, g and h).

Using the adult null model we quantified the percentile rank of each granule cell’s fraction of VGluT1-positive MF inputs relative to random sampling. Granule cells whose real fraction exceeded the 90th percentile of their null distribution were classified as VGluT1-positive-preferring (Fig. 2 i). Conversely, applying the same procedure to VGluT1-negative MFs identified VGluT1-negative-preferring granule cells (Fig. 2 j). Notably, cells that ranked in the top decile for VGluT1-positive input preference consistently fell in the bottom decile for VGluT1-negative preference, and vice versa (Fig. 2 I and j), reinforcing the existence of structured, non-random sampling subtypes. Based on these classifications, we divided the population into three groups: VGluT1-positive-preferring (n = 11), VGluT1-negative-preferring (n = 2), and non-preferring/neutral (n = 40) granule cells (see Fig. 2 k for their spatial distribution).

### Mossy Fiber-Centered Analysis Suggests Random Distribution of Output

We next examined mossy fiber terminal output patterns using a complementary null model approach. The first model (Fig. 3, b) simulated the adult state by constructing a sphere centered at each mossy fiber terminal using the average dendrite length (17.62 μm), and randomly assigned connections to granule cells whose centers of mass fell within the sphere. The number of connections was fixed to match the empirical data. The second model (Fig. 3, c) simulated a developmental state by using the maximum dendrite length (40.08 μm). For both models, we ran 1,000 Monte Carlo simulations.

**Figure 3.**
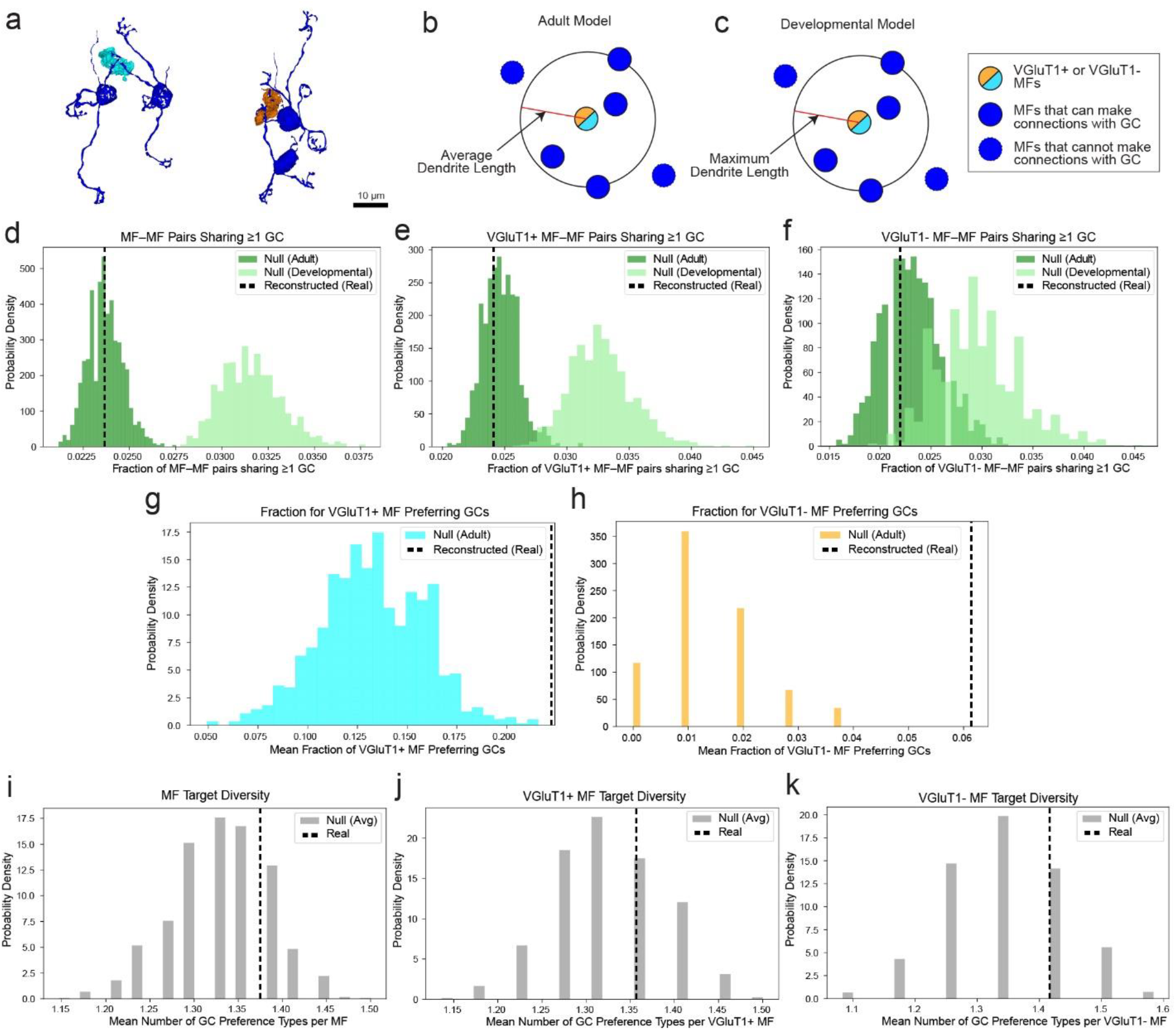
Mossy fibers distribute outputs in a largely random manner across granule cells. (**a**) 3D reconstructions of example mossy fiber (MF) terminals: a VGluT1+ terminal (cyan) contacting two granule cells (GCs, blue) and a VGluT1– terminal (orange) contacting two GCs. (**b, c**) Null models for MF-centered connectivity. (**b**) Adult null model: connections assigned within a sphere defined by the average dendrite length (17.62 μm). (**c**) Developmental null model: connections assigned within a sphere defined by the maximum dendrite length (40.08 μm). (**d–f**) Probability density distributions of the fraction of MF–MF pairs sharing ≥1 GC. Real data (black dashed line) are compared with null distributions from the adult (dark green) and developmental (light green) models. (**d**) All MFs, (**e**) only VGluT1+ MFs, (**f**) only VGluT1– MFs. (**g**,**h**) Probability density distributions of the mean fractions of VGluT1-positive-preferring (**g**) or VGluT1-negative-preferring (**h**) GC subtypes contacted by each MF in the real dataset (black dashed line) compared to the adult null model. (**i-k**) Probability density distributions of the mean target diversity, defined as the number of GC preference types (VGluT1-positive-preferring, VGluT1-negative-preferring, or non-preferring) contacted by each MF. (**i**) All MFs, (**j**) only VGluT1+ MFs, (**k**) only VGluT1– MFs.

We compared the fraction of mossy fiber pairs that shared at least one granule cell target in the empirical data to those in the null models. The observed overlap was not significantly different from the adult null model (Fig. 3, d; Adult null model, p-value (real < null) = 0.5340, Cohen’s d = -0.032) but was significantly lower than the developmental model (Fig. 3, d; Developmental null model, p-value (real < null) = 0.0000, Cohen’s d = -4.898). This indicates that mossy fibers distribute their outputs in a largely random manner in adulthood, with selective synapse making or pruning from development to adulthood likely contributing to the emergence of weak selectivity.

Breaking the analysis down by molecular identity, we found that VGluT1-positive terminals behaved similarly to the overall population (Fig. 3, e, VGluT1-positive MFs; Adult null model, p-value (real < null) = 0.3960, Cohen’s d = -0.374; Developmental null model, p-value (real < null) = 0.0000, Cohen’s d = -3.413), while VGluT1-negative terminals showed a weaker deviation from either adult or developmental null model (Fig. 3, f, VGluT1-negative MFs; Adult null model, p-value (real < null) = 0.4130, Cohen’s d = -0.398; Developmental null model, p-value (real < null) = 0.0150, Cohen’s d = -1.897). This suggests that VGluT1-negative terminals maintain a more random distribution pattern from VGluT1-positive terminals from development to adulthood, potentially as a strategy to ensure broad access by granule cells to these sparser inputs.

We further evaluated whether mossy fibers preferentially target specific granule cell subtypes as defined in the previous section. We compared the fraction of VGluT1-positive-preferring (Fig. 3, g) or VGluT1-negative-preferring (Fig. 3, h) granule cells contacted by each mossy fiber terminal to the adult null model. The observed values were significantly higher than expected by chance (Fig. 3, g and h; For VGluT1-positive-preferring GC fraction, p-value (real > null) = 0.0000; For VGluT1-negative-preferring GC fraction, p-value (real > null) = 0.0000), indicating that mossy fibers do bias connectivity toward granule cells predisposed to sample them.

Finally, we assessed the target diversity of granule cell targets across terminals by classifying GCs into three types: VGluT1-positive-preferring, VGluT1-negative-preferring, or others with the criterion of the top and bottom 10%. The target diversity value is defined by how many subtypes of GC contacted by a MF (1, 2, or 3). For all terminals, including VGluT1-positive and VGluT1-negative subsets, we found that the diversity of GC target types did not significantly differ from the null model (Fig. 3, i-k; All MF target diversity, p-value (real > null) = 0.238; VGluT1-positive MF target diversity, p-value (real > null) = 0.399; VGluT1-negative MF target diversity, p-value (real > null) = 0.341). This supports the idea that, while some degree of preferential targeting exists, mossy fiber terminals generally adopt a strategy to maximize distribution across available granule cells.

## Discussion

Our findings provide new insight into the structural logic of granule cell connectivity within the cerebellar cortex, demonstrating that granule cells sample mossy fiber inputs in a non-random, structured manner. Using molecularly defined terminals and spatially constrained null models, we reveal that granule cells exhibit input preferences that are stronger than expected by chance, particularly with respect to VGluT1-negative terminals, which are less abundant in this region (Han et al. 2024). This suggests that granule cells may compensate for input sparsity by adopting a broader sampling strategy for less common input types.

The contrast between granule cell–centered and mossy fiber–centered analyses highlights an important asymmetry in circuit organization. While granule cells appear selective in their sampling, mossy fiber terminals distribute their outputs in a manner largely consistent with random targeting. This discrepancy implies that input selection may be driven more by granule cell properties, such as dendritic reach or developmental pruning, rather than by targeted output from mossy fibers (Altman and Bayer 1978; Komuro and Rakić 1995; M. Kim et al. 2023; Schuldiner and Yaron 2015; Lackey, Heck, and Sillitoe 2018; Komuro and Rakic 1998). This also corresponds to the fact during development, mossy fiber enter the granule layer and form terminals before granule cells descend into granule layer (Schilling, Schmidt, and Baader 1994; Hámori and Somogyi 1983; T. Kim et al. 2023; Altman 1972). So it’s the granule cells that drive the selective synapse making.

Our modeling of both adult and developmental connectivity further supports a developmental refinement process. Patterns observed in the adult data deviate more strongly from chance than those in the developmental simulations, consistent with a pruning or competitive synapse making mechanism that enhances input diversification among neighboring granule cells (Hámori and Somogyi 1983; Lackey, Heck, and Sillitoe 2018; M. Kim et al. 2023; Dhar, Hantman, and Nishiyama 2018). The structured sampling of VGluT1-negative terminals may reflect a conserved strategy to ensure adequate integration of underrepresented input streams (Gebre, Reeber, and Sillitoe 2012; Nunzi, Russo, and Mugnaini 2003).

Recent connectomics studies using vEM have also demonstrated that granule cell input sampling is not entirely random. For example, granule cells have been reported to share mossy fiber partners more frequently than predicted by spatially constrained random models, and individual mossy fibers are unevenly represented across the granule cell population (Nguyen et al. 2022). In contrast, our analysis revealed the opposite trend: granule cells in our vCLEM dataset shared mossy fiber partners less often than expected under equivalent spatially constrained null models. This discrepancy may reflect differences in the cerebellar regions examined (vermis in (Nguyen et al. 2022) vs. Crus I in our study). Moreover, the underrepresented and overrepresented mossy fiber terminals described by (Nguyen et al. 2022) may correspond to distinct molecular subtypes, a distinction that was not accessible in their dataset because molecular identity was not labeled. By incorporating molecular identity through vCLEM, we extend these findings and demonstrate that structured sampling is especially evident for VGluT1-negative inputs. These results align with theoretical predictions that structured synaptic connectivity in the granule cell layer enhances pattern separation and expansion coding (Litwin-Kumar et al. 2017; Cayco-Gajic, Clopath, and Silver 2017).

The presence of granule cell subtypes with distinct input preferences suggests a potential basis for functional heterogeneity within the granule cell layer (Masoli et al. 2020; De Zeeuw, Lisberger, and Raymond 2021). These subtypes may contribute to parallel processing streams (Ishikawa, Shimuta, and Häusser 2015) or encode different aspects of sensorimotor information (Knogler et al. 2017; Sylvester et al. 2017), a hypothesis that warrants future investigation using other experimental approaches such as functional imaging or electrophysiology.

Our analysis was based on the published vCLEM dataset from the cerebellar Crus1 hemisphere, owing to the possibility of overlaying molecular identify on EM data with CLEM. To expand this type of analysis and deepen our understanding of how granule cells sample distinct mossy fiber inputs, generating more such correlative datasets on larger scales would be beneficial. Dye-based (Holtzman et al. 2009) or virus-based fluorescence tracing techniques (Pisano et al. 2021; Dohaku et al. 2019) can be combined to label mossy fibers from distinct regions. Furthermore, newer light microscopy based connectomics approach such like LICONN (Tavakoli et al. 2025) can help with accumulating more connectivity data with molecular identification.

Altogether, this study establishes a framework for examining how cerebellar microcircuits are wired to balance input molecular diversity, spatial constraints, and developmental process. Our findings open new avenues for understanding how structured sampling strategies support the integrative and computational functions of the cerebellum.

## Author Contribution

Xiaomeng Han: Conceptualization, data analysis, modeling, figure preparation, manuscript writing.

Elif Sevde Meral: Data segmentation, proofreading, data analysis, manuscript editing.

Jeff W. Lichtman: Supervision, conceptual input, manuscript editing, funding acquisition.

All authors read and approved the final manuscript.

## Acknowledgement

The vCLEM dataset used in this study was previously published (Han et al. 2024) and reused here with permission. We used OpenAI’s ChatGPT to assist with refining Python code and improving the clarity of the manuscript text. We thank Dr. Nagaraju Dhanyasi for helpful discussions on the manuscript. This work was supported by NIH R01 MH132710 (J. W. L., X. H.). E. S. M was supported by The Scientific and Technological Research Council of Turkiye (TUBITAK) 2209-A program, and mentored by Prof. Savaş Üstünova.

