## Supplementary Materials for "Structured Sampling of Molecularly Classified Mossy Fiber Inputs by Cerebellar Granule Cells"

Xiaomeng Han^1*^, Elif Sevde Meral^2^, Jeff Lichtman^1*^

1 Department of Molecular and Cellular Biology, Harvard University, Cambridge, MA

2 Bezmialem Vakif University School of Medicine, Istanbul, Turkey

* Corresponding authors

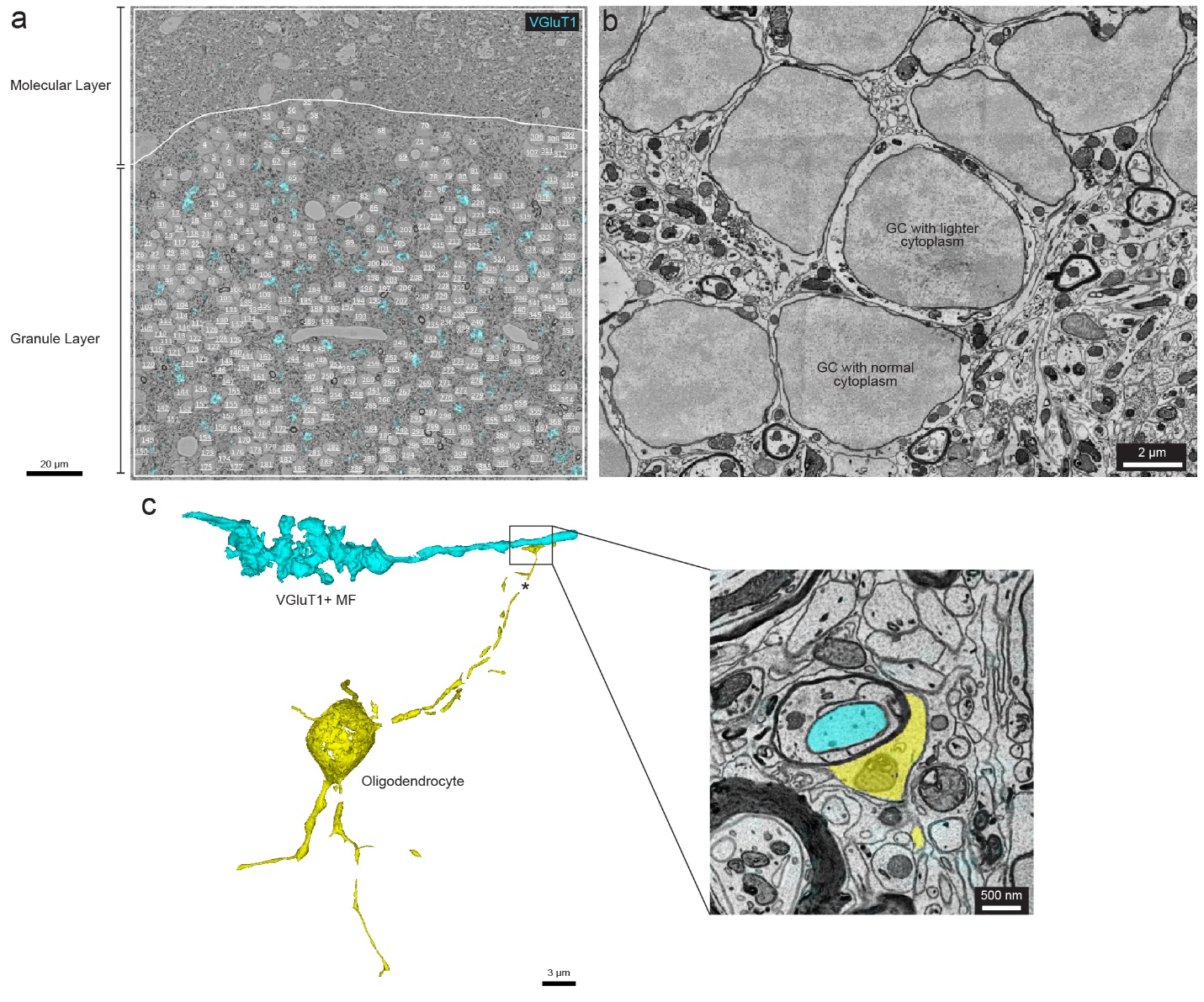

**Sup. Figure 1. Additional features of the granule cell layer in the vCLEM dataset.**

(**a**) Locations of 382 reconstructed cells (white labels) within the middle plane of the volume.

(**b**) EM micrograph showing examples of granule cells with normal versus lighter cytoplasm.

(**c**) 3D reconstruction of an oligodendrocyte (yellow) myelinating a VGluT1+ MF terminal (cyan). Right: EM inset of the myelination site. Asterisk marks unlabeled regions of the automatically segmented cell.
